# To cull or not to cull: a model-based evaluation of response strategies against Lumpy Skin Disease outbreaks

**DOI:** 10.64898/2025.12.31.697163

**Authors:** Thibaut Jombart, Jessica L. Abbate

## Abstract

With high risk of sustained transmission and substantial associated morbidity, Lumpy Skin Disease (LSD) of cattle presents a major emerging threat to bovine herds worldwide, and has the potential to cause huge losses for farmers, producers, and the wider bovine and dairy trade industries. The epidemic which started in June 2025 in France illustrates this threat, as the government-imposed mass culling strategy has led to protests by farmers and social unrest throughout the country. Here, we developed an infectious disease modelling framework for evaluating different intervention strategies in an effort to critically evaluate the evidence behind the drastic measures being taken. By simulating epidemics under different response settings, we found that control strategies employing targeted culling were essentially as effective as those requiring mass culling, and did not require significantly more vaccination. These results question the response strategy currently imposed by the French government, and suggest avenues for alternative response strategies which could gain better support from the affected communities. Our results are entirely reproducible using a new free, open-source simulation tool which can be used for further evaluation of public health interventions against LSD epidemics.

## INTRODUCTION

Lumpy skin disease (LSD) of cattle and related bovids is a vector-borne viral disease caused by lumpy skin disease virus (LSDV), a member of the genus *Capripoxvirus* in the family *Poxviridae*. The disease derives its name from the skin lesions that typically occur in the early stages of the disease and permanently scar the animals’ hides, often accompanied by fever, naso-pharyngeal secretions and swollen lymph nodes. While rarely fatal, LSDV can cause severe disease including pneumonia, lethargy, anorexia, emaciation, infertility and abortion, and reduced milk production that can last for several months ^1–4^.

The disease was first recognized in Zambia in 1929, where it was serologically and genetically nearly indistinguishable from sheep and goat pox viruses from the region, before spreading across the rest of the continent during the twentieth century ^1^. Since the 2010s, LSDV has emerged beyond Africa, with sustained transmission reported across the Middle East, large parts of Asia, and more recently has emerged into eastern and south-eastern Europe ^5,6^. Local transmission of LSDV is primarily thought to occur through the mechanical transfer of virus by blood-feeding insects, including biting flies (*Stomoxys* spp., in particular), certain mosquitoes, and hard ticks, rather than through direct animal-to-animal contact ^7,8^. Early observations noted that outbreaks tended to follow warm seasonal rains, when vector populations grew ^1,4^, a trend which followed its spread into Europe ^5,9^. However, long-distance spread is predominantly driven by human-mediated movement of infected cattle, often before clinical signs are apparent, leading to the seeding of new outbreak foci far beyond the range of vector dispersal, making it a worldwide threat ^10^.

Given the substantial economic burdens on cattle production, including extensive impacts on milk yield, decreased growth and weight gain, hide damage, and reproduction, new LSD outbreaks trigger emergency responses from agricultural authorities. Control of LSD outbreaks has relied on a combination of animal movement restrictions, vaccination, culling of infected animals, and vector control, though the relative emphasis placed on these measures has varied substantially between regions and epidemic contexts. In particular, mass vaccination with live attenuated capripoxvirus vaccines has been central to successful control in the Balkans and southeastern Europe, while culling has generally played a secondary role ^9,11,12^. The choice of control strategy is influenced by multiple contextual factors, including vector seasonality and abundance, cattle density and movement patterns, and concerns regarding adverse events or trade implications associated with the live vaccines currently available which render it impossible to distinguish vaccinated from naturally-infected animals. This last point is particularly important, as it raises fears of longer-term problems for regaining and retaining LSD-free trade status ^3^.

On 29 June 2025, France detected its first case of LSDV in the eastern department of Savoie, on the border with Italy which also confirmed its first outbreaks in 2025 ^13^. This development has sparked calls for coordination in deciding the best course of action to contain the emerging agricultural threat. The French government, in addition to preventative (ring) vaccination, restrictions on animal movements and increased sanitation protocols, adopted a policy of whole-herd culling including herds in which some animals had already been vaccinated. This approach provoked strong opposition from cattle farmers, who questioned the epidemiological necessity of blanket culling given the documented effectiveness of vaccination in other European settings ^5^. In December 2025, a new set of culling orders following spread to the southern region of Occitanie triggered protests, bringing broader public attention to the scientific and policy trade-offs underlying LSDV control ^14^. Both local and global stakes are high. According to figures from 2020, France is the 6th largest exporter of agricultural food products in the world, and milk and cattle products account for just over half of that production ^15^. Payments to just 42 affected livestock owners in Savoie totaled €2.4 million ^13^.

Given the economic and social impacts and associated tensions, the need for clear, well-reasoned, and evidence-driven LSD epidemic control decisions in the French context is of great importance. Here, we developed a spatially-explicit LSD compartmental transmission model, parameterized as much as possible to meet the realities of French surveillance capacities and response times, and designed to test the relative theoretical merits of different mixed-measure control strategies. We evaluate a range of possible response scenarios and compare the outcomes in terms of numbers of cases, deaths, geographic spread, and disease immunity. We illustrate how such models can be used for supporting decision-making and for communicating to the public the risks and benefits of those decisions, which is important for program compliance and success.

## MATERIAL AND METHODS

### Transmission model

We use a meta-population, stochastic compartmental model to capture the dynamics of LSD transmission. The model tracks the number of cattle in any of the following states at every point in time: susceptible (***S***), exposed (***E***), infectious (***I***), recovered (***R***), immunized through vaccination (***V***), dead from the disease (***D***), or culled (***C***). We also note ***N*** the total number of living individuals (***N*** = ***S*** + ***E*** + ***I*** + ***R*** + ***V***). Transitions between these states are summarised in Figure 1, and mathematical notations of the model detailed in Table 1. For convenience, we index time using ***t*** = **1**,**2, …, *T***, and population using ***j*** = **1**,**2, …, *J***, so that for instance, ***S***_***t,j***_ indicates the number of susceptible individuals at time ***t*** in patch ***j***.

**Table 1.** notations of the mathematical model of LSD transmission. Rates are indicated using Greek symbols, while proportions or probabilities are denoted using Latin symbols.

| Notation | Description |
| --- | --- |
| $J$ | the total number of populations in the meta-population; we usually use the indexing $j = 1, \dots, J$ |
| $T$ | the total number of time steps in a simulation; we usually use the indexing $t = 1, \dots, T$ |
| $\delta_{i,j}$ | proportion of the force of infection generated in population $i$ directed towards patch $j$ |
| $R_0$ | Basic reproduction number, used for calculating $\beta$, defined as the average number of secondary cases caused by an infectious individual in a fully susceptible population. |
| $\beta$ | individual rate of infection from infectious individuals ($I$) in the absence of response |
| $\sigma$ | rate at which exposed individuals ($E$) become infectious ($I$), treated as the inverse of the mean latency period |
| $\gamma$ | rate at which infected individuals ($I$) cease to be infected, treated as the inverse of the mean duration of illness |
| $m$ | case fatality ratio, defined as the proportion of infected individuals who die from the disease |
| $q^i, q^o$ | quarantine efficacy, in terms of reduction of transmission inside the population ($q^i$), or outwards to neighbouring populations ($q^o$) |
| $\omega_{t,j}$ | individual rate of vaccination at time $t$ in population $j$ |
| $\xi_{t,j}$ | culling rate at time $t$ in population $j$ |

**Figure 1.**
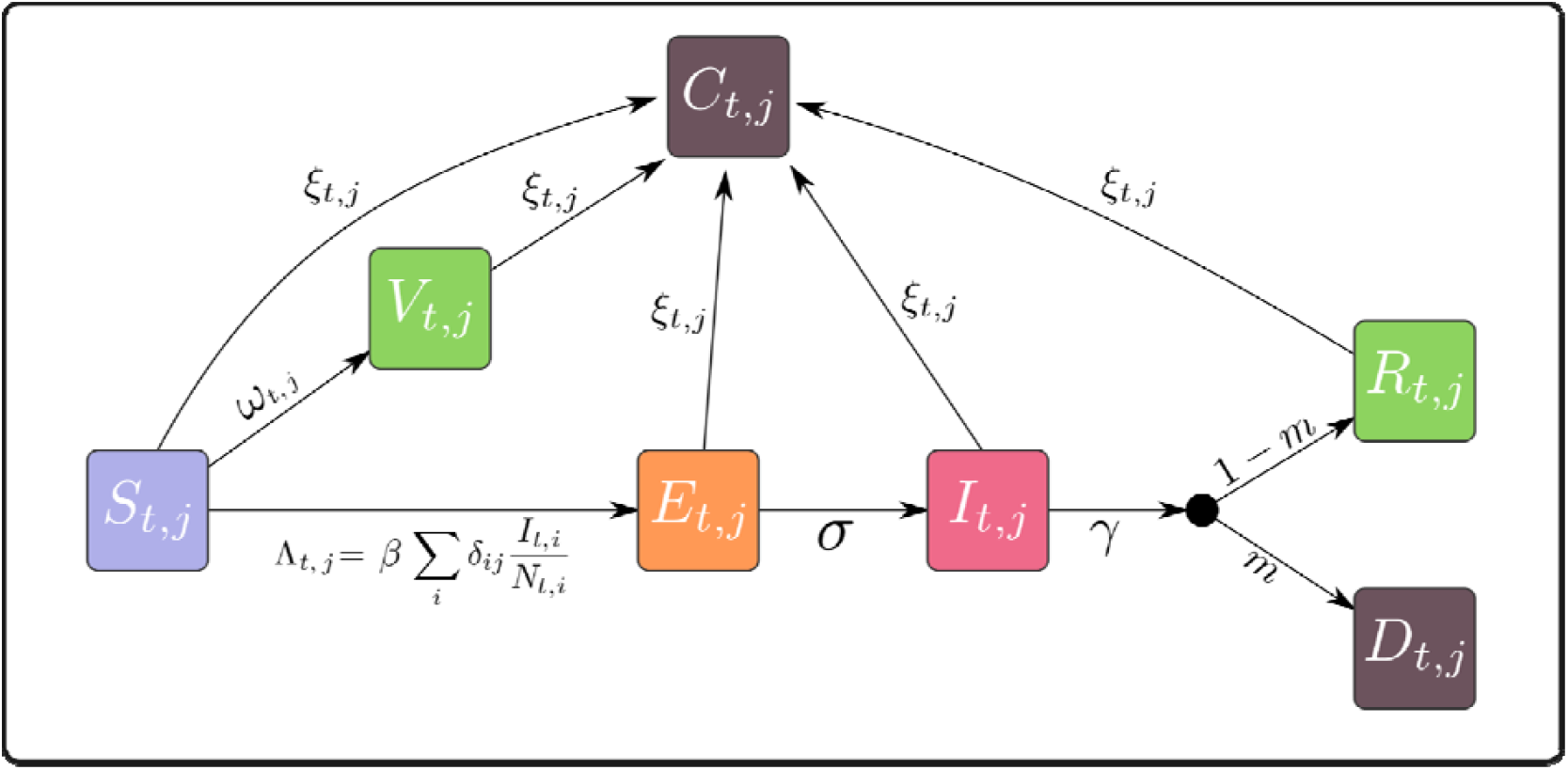
LSD transmission model. Each box represents a different state of disease dynamics including: susceptible (*S*), exposed (*E*), infectious (*I*), recovered (*R*), dead from the disease (*D*), vaccinated (*V*), or culled (*C*). Indices *t* and *j* representing time and space, respectively. These compartments are replicated for all populations, although a single one is represented for simplicity. Greek symbols indicate rates, while Latin symbols indicate probabilities. The black dot represents a bifurcation with two complementary probabilities (see Table 1).

Whenever a single transition is possible from state ***A*** to ***B*** and happens at a rate ***λ***, the number of individuals moving from ***A*** to ***B*** at time ***t*** in patch ***j***, denoted 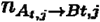, is drawn as:

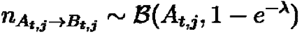

where **ℬ** indicates the Binomial distribution. In practice, most state changes in Figure 1 involve competing hazards, where individuals in a state ***A*** can transition to a number of states ***B***_**1**_, ***B***_**2**_, **…**, ***B***_***k***_ at rates ***λ***_**1**_, ***λ***_**2**_, **…**, ***λ***_***k***_. In such cases, we first draw the total number of individuals leaving ***A*** as:

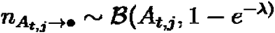

where 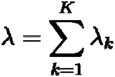. We then draw the numbers of transitions to the respective compartments as:

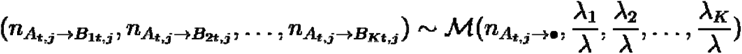

where **ℳ** indicates the Multinomial distribution.

These principles apply to all changes illustrated in Figure 1, for each population in the meta-population. Details of individual equations are provided in Text S1.

In our simulations (see next section), we assume populations are farms, arranged on a regular 2-dimensional grid, but our framework can accommodate any geographic structure. The connectivity between populations is modelled as a matrix **Δ** = **{*δ***_***i,j***_**}** where ***δ***_***i,j***_ is the proportion of the force of infection generated in population directed towards population ***j***. The diagonal terms ***δ***_***i,i***_ represent the proportion of the force of infection generated in creating new cases inside the same population. It follows that the force of infection experienced by population ***j*** as time ***t***, denoted **Λ**_***t,j***_, can be calculated as:

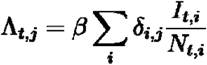

Note that this formulation enables interventions affecting transmissibility to be geographically targeted, as they can impact transmission within a specific patch ***i*** by reducing ***δ***_***i,i***_, or transmission towards ‘s neighbouring populations by reducing ***δ***_***i,j,i*≠ *j***_. This is used, for instance, to simulate quarantine and vector control (considered here as the use of insecticides; see Table 3).

The rates of vaccination ***ω***_***t,j***_ and culling ***ξ***_***t,j***_ are time- and space-dependent to enable the activation of specific interventions: they are normally 0 unless a response has been triggered in a given population, in which case these parameters can take non-null values (see Table 2 and 3).

**Table 2.** values of parameters used in LSD epidemics simulations in a meta-population of farms.

| Parameter | Value | Source |
| --- | --- | --- |
| $J$ | 1600 farms arranged on a two-dimensional regular grid (40x40) | - |
| $T$ | 365 days | - |
| $S_j$ | The herd size of farms was drawn as $20 + X$ where $X$ is exponentially distributed with mean 30, to replicate a long-tailed distribution with an average size of 50 cattle | 16 |
| $R_0$ | Log-normal distribution with mean 9.7 and standard deviation 0.05 | 20 |
| $\gamma$ | Log-normal distribution with mean 8.8 and standard deviation 0.05 | 20 |
| $\beta$ | Calculated as $R_0\gamma$ | 26 |
| $\sigma$ | Log-normal distribution with mean 8.1 and standard deviation 0.05 | 20 |
| $m$ | Log-normal distribution with mean 0.03 and standard deviation 0.2 | 27–29 |
| $\omega_{t,j}$ | When vaccination does occur, we assumed 100% vaccine coverage, with vaccine efficacy linearly increasing to 65% at 21 days post injection, resulting in a daily immunisation rate of $-\log(1 - 0.65/21)$ | 30 |
| $\xi_{t,j}$ | When culling does occur, we assumed a rate of 0.5, so that on average 50% of the herd is killed each day | - |

**Table 3.** implementation of response components. This table outlines how different response components were implemented in the simulations. Each simulation scenario is a combination of some of these components (see Table 4). ‘Notation’ corresponds to elements used in scenario labels. See Table 2 for explanations on the values of intervention parameters.

| Response component | Notation | Implementation |
| --- | --- | --- |
| Mass culling | *cull* | Individuals in the affected farm are killed at a rate $\xi = 0.5$ |
| Selective culling | *selcull* | Infectious individuals (*I*) in the affected farm are culled immediately. |
| Vaccination | *vacc* | Individuals in the affected farm and the neighbouring farms are vaccinated at a rate $\omega_{i,j} = -\log(1 - 0.65/21)$ |
| Quarantine | *quarant* | Reduction of transmission inside the affected farm by 20%.<br>Reduction of transmission towards the neighbouring farms by 90%. |
| Insecticide | *insect* | Insecticide is applied in the affected farm and the neighbouring farms, reducing transmission by 50% |

Note: the notion of ‘neighbouring’ farms uses the rook connectivity (see Figure S2). All responses take place 4 days after the first symptom onset (first *I*), and end 28 days after the last case.

### Epidemics and response simulations

We used our model to simulate epidemics of LSD in cattle herds on a network of inter-connected farms. Herd sizes were simulated using an inflated exponential distribution, with an average of 50 cattle per herd and a minimum size of 20 (Table 2, Figure S1), in line with reported herd size of cattle farms in France ^16^. Each simulation involved a set of 1,600 farms (each farm being treated as a ‘population’) organized on a 40x40 regular grid (Figure S2). Rook connectivity ^17^ was used to define neighbours, and the corresponding connectivity matrix **Δ** was defined so that 5% of transmission was uniformly directed towards neighbouring farms, with the remaining 95% of transmission happening within farms. We also seeded epidemics by adding one initial infectious individual (***I***) to a randomly selected farm at the first time step of each simulation. Taken together, herd sizes, their geographic distribution, and the location of the initial case form what we refer to as ‘initial conditions’.

A total of 1,000 initial conditions were generated randomly. For each, the transmission process defined by our model was simulated daily for 365 days, using different types of interventions (Table 3) combined into response scenarios (Table 4). All interventions took place 4 days after the first case (***I***) in an affected farm, and lasted for 28 days after the last case. Five types of interventions were then considered (Table 3): mass culling (labelled: ‘*cull*’) of herds with at least one infectious individual (***I***), selective culling (‘*selcull*’) where only infectious individuals are killed, quarantine (‘*quarant*’) which reduces transmission within the affected farm by 20% and towards other farms by 90%, mass vaccination (‘*vacc*’) of all individuals in the affected farm and the neighbouring farms, and insecticide use (‘*insect*’) in the affected farm and the neighbouring farms reducing overall transmission by 50%. These interventions were combined into 6 different response scenarios, ranging from a baseline scenario with no response, to scenarios combining some or all interventions (Table 6). Of note, the scenario labelled ‘*cull_quarant_vacc*’ is closest to the response currently in place in France. Each scenario was applied to each initial condition, so that a total of 6,000 simulations were produced and analysed.

**Table 4.** structure of simulated scenarios. This table outlines the different simulated response scenarios. All scenarios use the same set of 1,000 simulated meta-populations. See Table 3 for details on the implementation of the response components.

| Scenario label | Description |
| --- | --- |
| *baseline* | Baseline scenario with no response |
| *cull_quarant_vacc* | Mass culling of infected herds (with at least one *I*), quarantine of infected farms and neighbouring farms, and vaccination of herds in neighbouring farms |
| *quarant_vacc* | Quarantine of infected farms and neighbouring farms, and vaccination of herds in neighbouring farms |
| *selcull_quarant_vacc* | Like *quarant_vacc*, with the addition of selective culling of infectious individuals only (*I*) |
| *selcull_quarant_vacc_insect* | Like *selcull_quarant_vacc*, with the addition of insecticide use in affected farms and their neighbours |
| *all* | Like *all_but_cull* but mass culling replaces selective culling |

Note: the notion of 'neighbouring' farms uses the rook connectivity (see Figure S2). All responses take place 7 days after the first symptom onset (first *I*), and end 28 days after the last case.

### Data availability and reproducibility

The model as well as tools for generating meta-population structures and LSD transmission parameters are available in the new free, open-source package *lsdsim* for the R software ^18^, available on github at: https://github.com/thibautjombart/lsdsim

The scripts and infrastructure for running simulations and reproducing the results presented here are available as an instance of a *reportfactory* ^19^, available from the same github repository.

## RESULTS

Figure 2 provides a graphical summary of the results of our simulation study, and numeric details are provided in Tables S1-S4. Of first note, the baseline scenario, using no response at all, shows that the epidemic spreads widely, with tens of thousands of cases (Figure 2A) and hundreds of farms being infected (Figure 2B). In fact, in all of these simulations the epidemic was still spreading after a year, so that the final number of cases would likely be much higher if simulations carried on for longer. Because of the large number of infected cattle and the relatively low mortality, this scenario also resulted in non-negligible immunity in the population (Figure 2C). The absolute number of deaths remained high, however, due to the sheer number of cases, making it the worst case scenario in terms of overall loss of individuals (Figure 2D).

**Figure 2.**
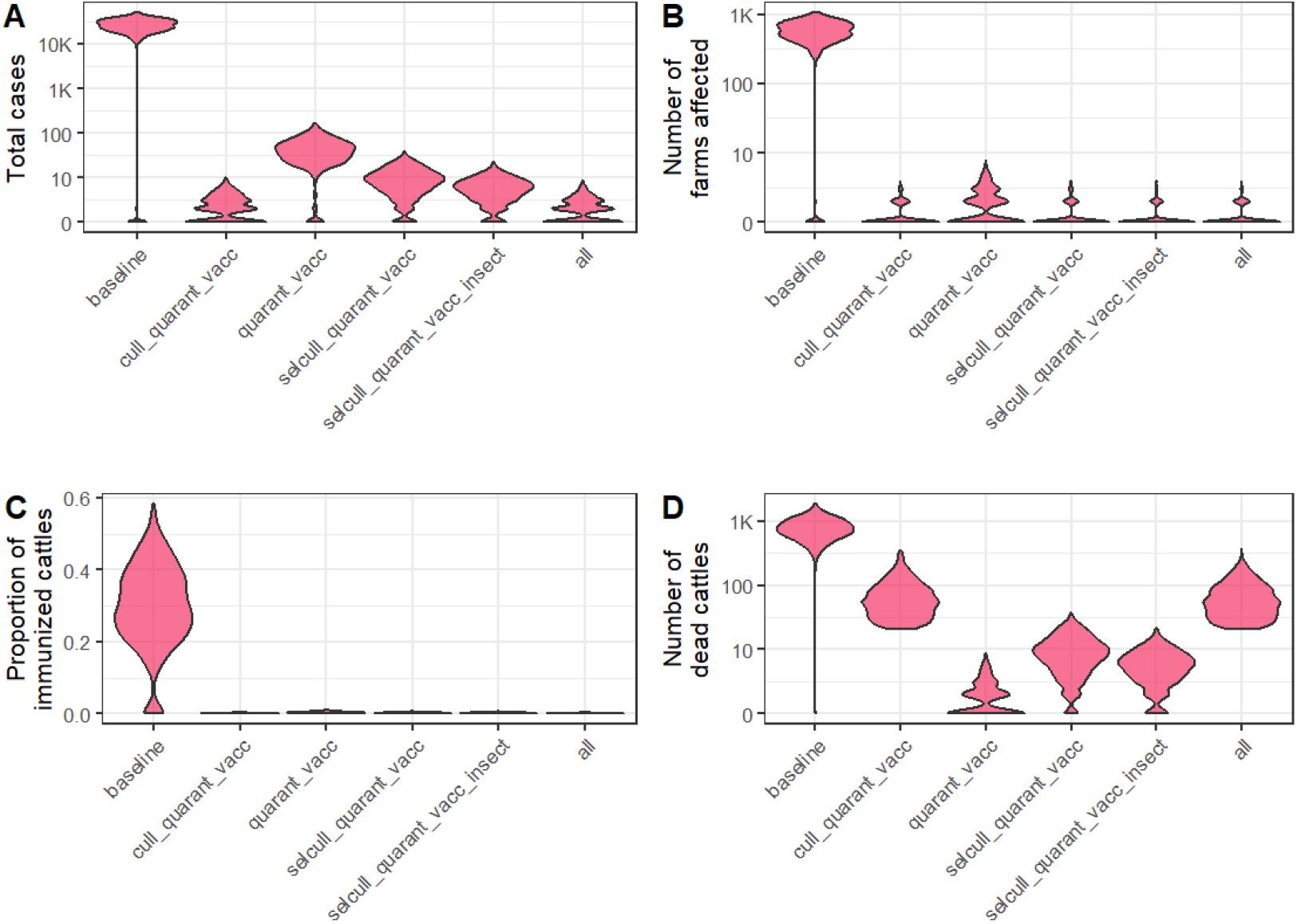
simulations of LSD epidemics under different response scenarios. Violinplots represent the density of simulations for different measured outcome and response scenarios (see Table 4). Each violin represents 1,000 simulations. Results are aggregated across all farms, 365 days after the first introduction of LSD in a randomly chosen farm. **A.** Total number of cases (including *E, I, R*, and *D*). **B**. Total number of farms having experienced at least one case. **C**. Proportion of immunized cattles, including *V* and *R*. **D**. Number of dead cattles, including *C* and *D*. Note that the y axis uses log-10 scales in panels A and B. Details of corresponding numbers can be found in Tables S1-4.

The currently ongoing response in France using a combination of mass culling, quarantine, and vaccination (‘*cull_quarant_vacc*’) achieved the best results in terms of minimizing the number of cases (Figure 2A, Tables S1, S3), but was also second worst in terms of overall deaths due to disease mortality and culling (Figure 2D, Table S2). It was otherwise indistinguishable from other interventions when considering geographic spread and overall immunity (Figure 2B). Indeed, all other scenarios involving different combinations of quarantine, vaccination, selective culling and insecticide use yielded similar results, with very limited numbers of cases and geographic spread, and very low numbers of deaths except where selective culling was used. All interventions led to negligible immunity in the population a year after the first LSD introduction (Figure 2C), likely owing to the small numbers of farms affected. A good compromise to limit both cases and deaths was achieved by a combination of selective culling, quarantine, and vaccination, which kept outbreak size to tens of cases while also minimizing the loss of individuals (Figure 2, Tables S1-S4). The addition of insecticide use further reduced the number of cases, with no noticeable impact on overall deaths.

## DISCUSSION

With the potential for very high and sustained inter-annual transmissibility, and considerable impacts on animal health and productivity, LSD is a major threat to cattle herds worldwide. Due to the high connectivity between remote herds due to national and international trade, introductions of LSD in countries where it is not yet endemic are going to be increasingly likely, and it is paramount to devise data-driven response strategies for outbreak management. Here, we have developed a spatialized, stochastic compartmental model to capture key aspects of LSD transmission. Using simulations, we have compared the impact of different response scenarios a year after an initial introduction of LSD in terms of numbers of cases, geographic spread, population-level immunity, and deaths.

LSD is highly contagious, with basic reproduction numbers which can exceed 10 depending on the insect vectors and season considered ^20^. Our findings confirm that in the absence of response, a single LSD case introduced in a farm can quickly lead to large outbreaks, first inside the herd, but also to neighbouring farms, ultimately leading to very high attack rates and large numbers of deaths in the entire meta-population. Our results also show that the strategy currently advocated by the French government, involving mass culling of herds with at least one infectious individual and vaccination of cattle in neighbouring farms, is likely to contain further spread. However, we also show that other interventions, involving only selective culling of infectious individuals, or even no culling at all, are essentially as likely to contain nascent epidemics, with lower overall death tolls and no major increase in the number of vaccinated (or naturally sero-converted) animals required.

These results are in line with the findings of prior modeling and responses in other European contexts, which found that mass culling was only necessary when vaccination coverage was too low ^5,9,11,21^. Of all considered alternatives, we posit that the combination of selective culling, quarantine, and ring vaccination offers a good compromise between limiting the number of cases and overall loss of individuals. Insecticide use would further reduce cases, but we argue that the impact on farmers and cattle’s health, on the environment, as well as potential for selecting resistance in the insect populations, should be considered before recommending this strategy.

The relative cost-effectiveness of the different strategies remain to be investigated. Indeed, the strongest economic impact of LSD epidemics arguably lies in the associated morbidity, rather than mortality alone ^2,3^. Combining a costing model for the different types of interventions listed in Table 3 to our simulation framework would yield further insights into optimal strategies, and permit to weigh appropriately the relative costs of LSD-related death and morbidity.

This study provides a proof-of-concept that alternative strategies may be more desirable than mass culling to respond to LSD outbreaks, while remaining just as effective, but our results will be improved by better parametrisation. Indeed, while we informed our model with literature-derived estimates for transmission parameters and the natural history of LSD, *ad-hoc* values were chosen for some parameters relating to the interventions, informed where possible by personal communications with farmers and veterinarians in France. Ideally, field studies, or polls from a representative panel of experts including farmers and veterinarians, should be used to improve the realism of our simulations. We also note that other interventions, or combinations of interventions into scenarios, could be considered.

This work uses a simplified meta-population setting which, while using realistic herd sizes, cannot be used for forecasting trajectories of actual epidemics. However, the flexibility of our framework readily enables simulations using actual herd sizes, geographic distribution of farms, and connectivity, which would yield much more realistic results. Further work should also focus on improving our understanding of the persistence of the disease in the environment, as well as repeated introductions through international trade, and the consequences of these phenomena on longer-term LSD dynamics. Such studies will be essential for assessing the scale and feasibility of vaccination strategies to prevent future re-emergence ^21^.

One last key aspect, not considered in our simulations, but which motivated this work and lies behind the large protests against the LSD response in France, is social acceptance. As illustrated during other infectious diseases epidemics which have had large social impacts ^22–25^, acceptance is key to ensuring an efficient response. Simply put, a response strategy which would be theoretically best at containing an outbreak, or most cost-effective, is only as good as it is feasible, and feasibility requires acceptance (facilitating cooperation and compliance) by the affected communities. By providing a theoretical framework and free, open-source simulation tool to evaluate the impact of different LSD responses, we hope this works contributes to opening up a rational debate on alternative LSD response strategies.

## ACKNOWLEDGEMENTS

TJ acknowledges funding from the MRC Centre for Global Infectious Disease Analysis (reference MR/X020258/1), funded by the UK Medical Research Council (MRC). This UK funded award is carried out in the framework of the Global Health EDCTP3 Joint Undertaking. We thank Dr Laurent Morvilliers and the unnamed farmers who helped inform the parametrisation of interventions used in our simulations. No generative AI was used in any part of this work.

## AUTHOR CONTRIBUTIONS

TJ developed and implemented the transmission model, ran simulations, generated tables and figures, and wrote the manuscript. JLA reviewed the literature, reviewed and discussed results interpretation, and wrote the manuscript.

