## Supplementary Material for "To cull or not to cull: a model-based evaluation of response strategies against Lumpy Skin Disease outbreaks"

**Table of contents**

**Text S1: Details of model transitions.	2**

**Figure S1: Herd size distribution used in simulated LSD epidemics.	3**

**Figure S2: Example of geographic distribution of farms.	4**

**Table S1: Number of LSD cases in the simulated epidemics.	5**

**Table S2: Number of LSD deaths in the simulated epidemics.	6**

**Table S3: Attack rates of LSD in the simulated epidemics.	7**

**Table S4: Number of farms with at least one LSD case in the simulated epidemics.	8**

### Text S1: Details of model transitions.

This section details the equations corresponding to the transitions illustrated in Figure 1. Competing hazards are handled using a two-step process: first drawing the total number of individuals leaving a compartment using a Binomial draw, and then assigning new states using a Multinomial distribution.

Globally, the system is described by the set of equations:

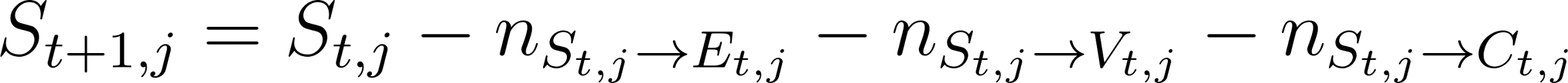
<https://www.codecogs.com/eqnedit.php?latex=%20S_%7Bt%2B1%2Cj%7D%20%3D%20S_%7Bt%2Cj%7D%20-%20n_%7BS_%7Bt%2Cj%7D%20%5Crightarrow%20E_%7Bt%2Cj%7D%7D%20-%20n_%7BS_%7Bt%2Cj%7D%20%5Crightarrow%20V_%7Bt%2Cj%7D%7D%20-%20n_%7BS_%7Bt%2Cj%7D%20%5Crightarrow%20C_%7Bt%2Cj%7D%7D%20#0>

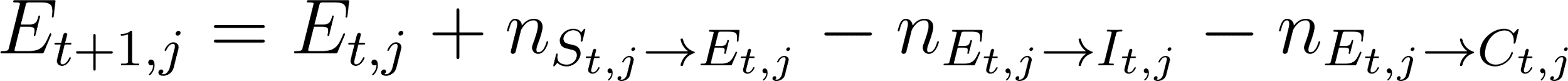
<https://www.codecogs.com/eqnedit.php?latex=%20E_%7Bt%2B1%2Cj%7D%20%3D%20E_%7Bt%2Cj%7D%20%2B%20n_%7BS_%7Bt%2Cj%7D%20%5Crightarrow%20E_%7Bt%2Cj%7D%7D%20-%20n_%7BE_%7Bt%2Cj%7D%20%5Crightarrow%20I_%7Bt%2Cj%7D%7D%20-%20n_%7BE_%7Bt%2Cj%7D%20%5Crightarrow%20C_%7Bt%2Cj%7D%7D%20#0>

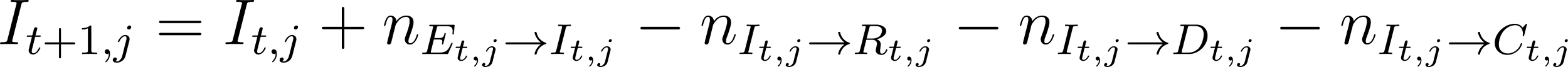
<https://www.codecogs.com/eqnedit.php?latex=%20I_%7Bt%2B1%2Cj%7D%20%3D%20I_%7Bt%2Cj%7D%20%2B%20n_%7BE_%7Bt%2Cj%7D%20%5Crightarrow%20I_%7Bt%2Cj%7D%7D%20-%20n_%7BI_%7Bt%2Cj%7D%20%5Crightarrow%20R_%7Bt%2Cj%7D%7D%20-%20n_%7BI_%7Bt%2Cj%7D%20%5Crightarrow%20D_%7Bt%2Cj%7D%7D%20%20-%20n_%7BI_%7Bt%2Cj%7D%20%5Crightarrow%20C_%7Bt%2Cj%7D%7D%20#0>

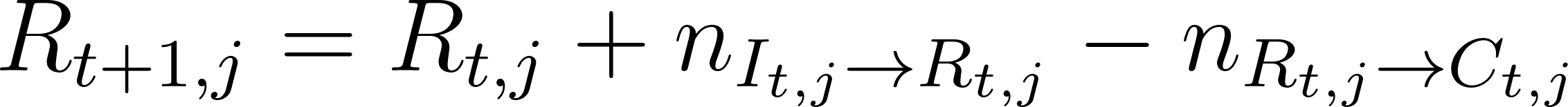
<https://www.codecogs.com/eqnedit.php?latex=%20R_%7Bt%2B1%2Cj%7D%20%3D%20R_%7Bt%2Cj%7D%20%2B%20n_%7BI_%7Bt%2Cj%7D%20%5Crightarrow%20R_%7Bt%2Cj%7D%7D%20-%20n_%7BR_%7Bt%2Cj%7D%20%5Crightarrow%20C_%7Bt%2Cj%7D%7D%20#0>

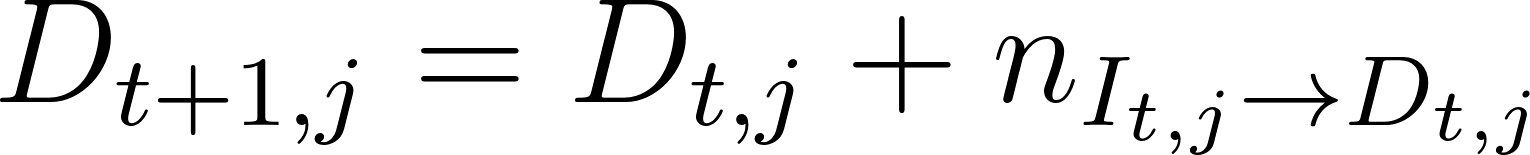
<https://www.codecogs.com/eqnedit.php?latex=%20D_%7Bt%2B1%2Cj%7D%20%3D%20D_%7Bt%2Cj%7D%20%2B%20n_%7BI_%7Bt%2Cj%7D%20%5Crightarrow%20D_%7Bt%2Cj%7D%7D%20#0>

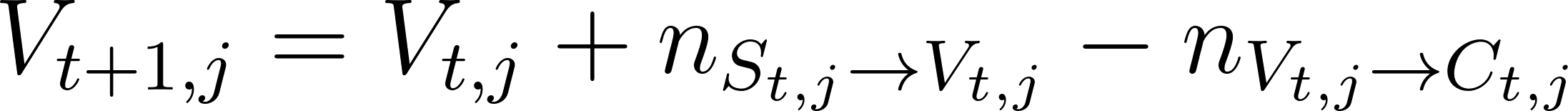
<https://www.codecogs.com/eqnedit.php?latex=%20V_%7Bt%2B1%2Cj%7D%20%3D%20V_%7Bt%2Cj%7D%20%2B%20n_%7BS_%7Bt%2Cj%7D%20%5Crightarrow%20V_%7Bt%2Cj%7D%7D%20-%20n_%7BV_%7Bt%2Cj%7D%20%5Crightarrow%20C_%7Bt%2Cj%7D%7D%20#0>

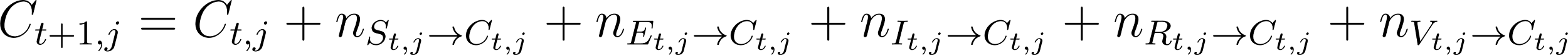
<https://www.codecogs.com/eqnedit.php?latex=%20C_%7Bt%2B1%2Cj%7D%20%3D%20C_%7Bt%2Cj%7D%20%2B%20n_%7BS_%7Bt%2Cj%7D%20%5Crightarrow%20C_%7Bt%2Cj%7D%7D%20%2B%20n_%7BE_%7Bt%2Cj%7D%20%5Crightarrow%20C_%7Bt%2Cj%7D%7D%20%2B%20n_%7BI_%7Bt%2Cj%7D%20%5Crightarrow%20C_%7Bt%2Cj%7D%7D%20%2B%20n_%7BR_%7Bt%2Cj%7D%20%5Crightarrow%20C_%7Bt%2Cj%7D%7D%20%2B%20n_%7BV_%7Bt%2Cj%7D%20%5Crightarrow%20C_%7Bt%2Cj%7D%7D#0>

The number of individuals leaving
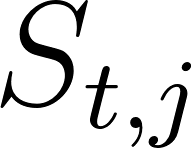
<https://www.codecogs.com/eqnedit.php?latex=S_%7Bt%2Cj%7D#0> is determined as:

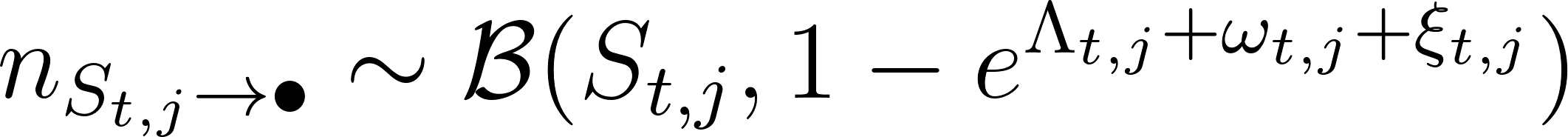
<https://www.codecogs.com/eqnedit.php?latex=%20n_%7BS_%7Bt%2Cj%7D%20%5Crightarrow%20%5Cbullet%7D%20%5Csim%20%5Cmathcal%7BB%7D%20(S_%7Bt%2Cj%7D%2C%201%20-%20e%5E%7B%5CLambda_%7Bt%2Cj%7D%20%2B%20%5Comega_%7Bt%2Cj%7D%20%2B%20%5Cxi_%7Bt%2Cj%7D%7D)%20#0>

Then the number of individuals going from
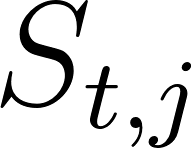
<https://www.codecogs.com/eqnedit.php?latex=S_%7Bt%2Cj%7D#0> to
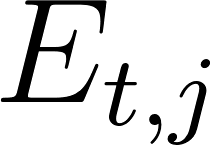
<https://www.codecogs.com/eqnedit.php?latex=E_%7Bt%2Cj%7D#0>,
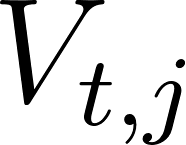
<https://www.codecogs.com/eqnedit.php?latex=V_%7Bt%2Cj%7D#0> and
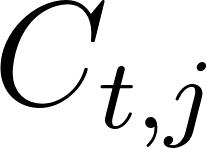
<https://www.codecogs.com/eqnedit.php?latex=C_%7Bt%2Cj%7D#0> is determined as:

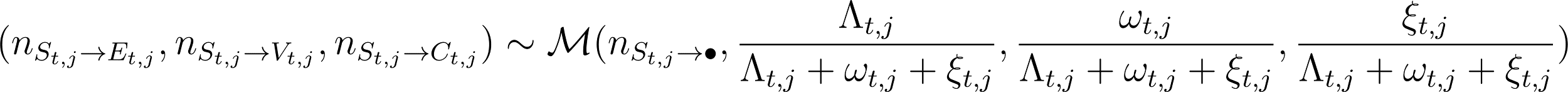
<https://www.codecogs.com/eqnedit.php?latex=%20(n_%7BS_%7Bt%2Cj%7D%20%5Crightarrow%20E_%7Bt%2Cj%7D%7D%2C%20n_%7BS_%7Bt%2Cj%7D%20%5Crightarrow%20V_%7Bt%2Cj%7D%7D%2C%20n_%7BS_%7Bt%2Cj%7D%20%5Crightarrow%20C_%7Bt%2Cj%7D%7D)%20%5Csim%20%5Cmathcal%7BM%7D(n_%7BS_%7Bt%2Cj%7D%20%5Crightarrow%20%5Cbullet%7D%2C%20%5Cfrac%7B%5CLambda_%7Bt%2Cj%7D%7D%7B%5CLambda_%7Bt%2Cj%7D%20%2B%20%5Comega_%7Bt%2Cj%7D%20%2B%20%5Cxi_%7Bt%2Cj%7D%7D%2C%20%20%5Cfrac%7B%5Comega_%7Bt%2Cj%7D%7D%7B%5CLambda_%7Bt%2Cj%7D%20%2B%20%5Comega_%7Bt%2Cj%7D%20%2B%20%5Cxi_%7Bt%2Cj%7D%7D%2C%20%5Cfrac%7B%5Cxi_%7Bt%2Cj%7D%7D%7B%5CLambda_%7Bt%2Cj%7D%20%2B%20%5Comega_%7Bt%2Cj%7D%20%2B%20%5Cxi_%7Bt%2Cj%7D%7D)#0>

The number of individuals leaving
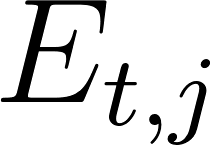
<https://www.codecogs.com/eqnedit.php?latex=E_%7Bt%2Cj%7D#0> is determined as:

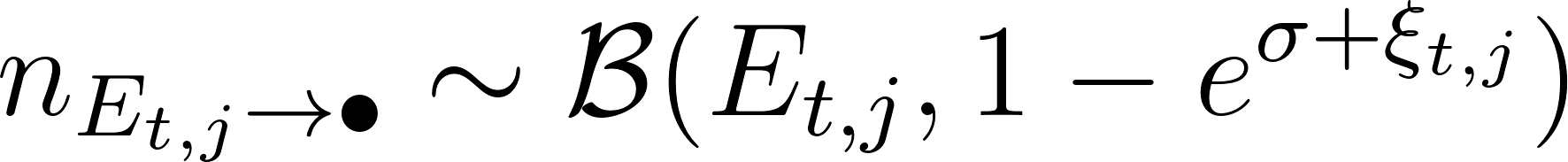
<https://www.codecogs.com/eqnedit.php?latex=%20n_%7BE_%7Bt%2Cj%7D%20%5Crightarrow%20%5Cbullet%7D%20%5Csim%20%5Cmathcal%7BB%7D%20(E_%7Bt%2Cj%7D%2C%201%20-%20e%5E%7B%5Csigma%20%2B%20%5Cxi_%7Bt%2Cj%7D%7D)%20#0>

Then the number of individuals going from
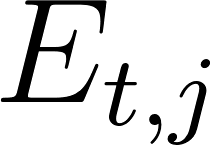
<https://www.codecogs.com/eqnedit.php?latex=E_%7Bt%2Cj%7D#0> to
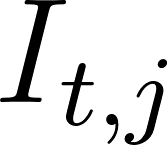
<https://www.codecogs.com/eqnedit.php?latex=I_%7Bt%2Cj%7D#0> and
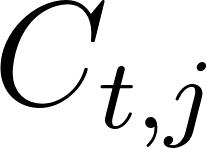
<https://www.codecogs.com/eqnedit.php?latex=C_%7Bt%2Cj%7D#0> is determined as:

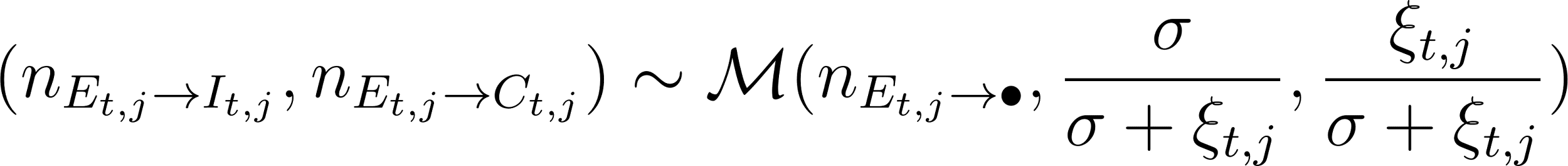
<https://www.codecogs.com/eqnedit.php?latex=%20(n_%7BE_%7Bt%2Cj%7D%20%5Crightarrow%20I_%7Bt%2Cj%7D%7D%2C%20n_%7BE_%7Bt%2Cj%7D%20%5Crightarrow%20C_%7Bt%2Cj%7D%7D)%20%5Csim%20%5Cmathcal%7BM%7D(n_%7BE_%7Bt%2Cj%7D%20%5Crightarrow%20%5Cbullet%7D%2C%20%5Cfrac%7B%5Csigma%7D%7B%5Csigma%20%2B%20%5Cxi_%7Bt%2Cj%7D%7D%2C%20%20%5Cfrac%7B%5Cxi_%7Bt%2Cj%7D%7D%7B%5Csigma%20%2B%20%5Cxi_%7Bt%2Cj%7D%7D)#0>

The number of individuals leaving
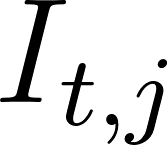
<https://www.codecogs.com/eqnedit.php?latex=I_%7Bt%2Cj%7D#0> is determined as:

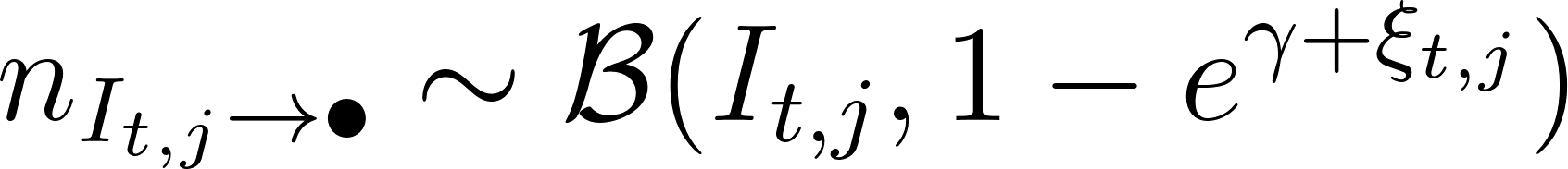
<https://www.codecogs.com/eqnedit.php?latex=%20n_%7BI_%7Bt%2Cj%7D%20%5Crightarrow%20%5Cbullet%7D%20%5Csim%20%5Cmathcal%7BB%7D%20(I_%7Bt%2Cj%7D%2C%201%20-%20e%5E%7B%5Cgamma%20%2B%20%5Cxi_%7Bt%2Cj%7D%7D)%20#0>

Then the number of individuals going from
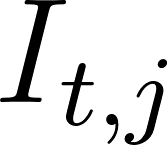
<https://www.codecogs.com/eqnedit.php?latex=I_%7Bt%2Cj%7D#0> to
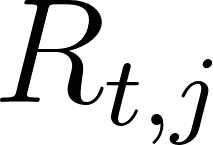
<https://www.codecogs.com/eqnedit.php?latex=R_%7Bt%2Cj%7D#0>,
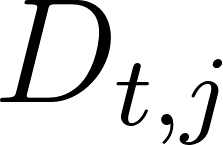
<https://www.codecogs.com/eqnedit.php?latex=D_%7Bt%2Cj%7D#0> and
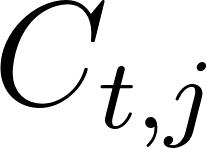
<https://www.codecogs.com/eqnedit.php?latex=C_%7Bt%2Cj%7D#0> is determined using:

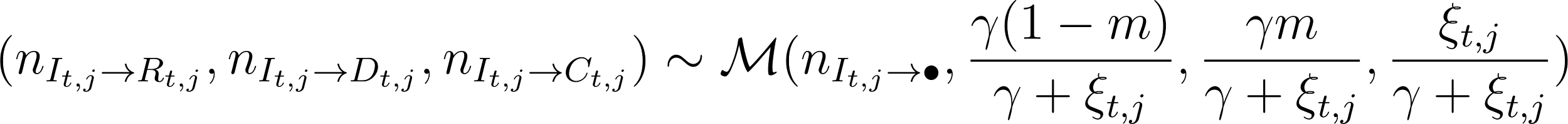
<https://www.codecogs.com/eqnedit.php?latex=%20(n_%7BI_%7Bt%2Cj%7D%20%5Crightarrow%20R_%7Bt%2Cj%7D%7D%2C%20n_%7BI_%7Bt%2Cj%7D%20%5Crightarrow%20D_%7Bt%2Cj%7D%7D%2C%20n_%7BI_%7Bt%2Cj%7D%20%5Crightarrow%20C_%7Bt%2Cj%7D%7D)%20%5Csim%20%5Cmathcal%7BM%7D(n_%7BI_%7Bt%2Cj%7D%20%5Crightarrow%20%5Cbullet%7D%2C%20%5Cfrac%7B%5Cgamma%20(1-m)%7D%7B%5Cgamma%20%2B%20%5Cxi_%7Bt%2Cj%7D%7D%2C%20%5Cfrac%7B%5Cgamma%20m%7D%7B%5Cgamma%20%2B%20%5Cxi_%7Bt%2Cj%7D%7D%2C%20%5Cfrac%7B%5Cxi_%7Bt%2Cj%7D%7D%7B%5Cgamma%20%2B%20%5Cxi_%7Bt%2Cj%7D%7D%20)#0>

Finally, the number of individuals leaving
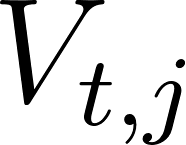
<https://www.codecogs.com/eqnedit.php?latex=V_%7Bt%2Cj%7D#0> and
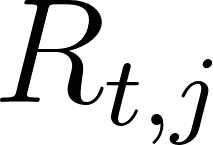
<https://www.codecogs.com/eqnedit.php?latex=R_%7Bt%2Cj%7D#0> determined as, respectively:

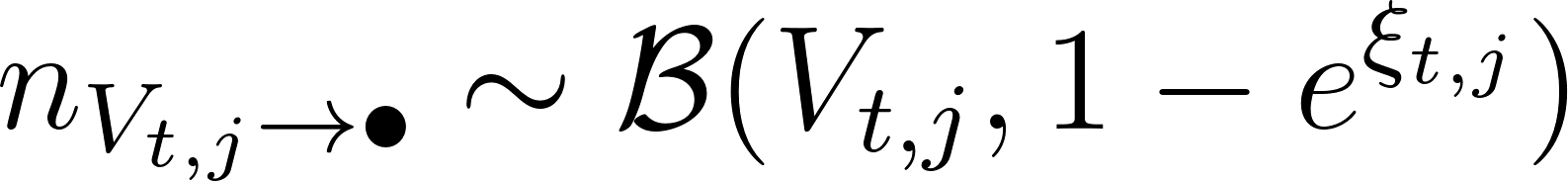
<https://www.codecogs.com/eqnedit.php?latex=%20n_%7BV_%7Bt%2Cj%7D%20%5Crightarrow%20%5Cbullet%7D%20%5Csim%20%5Cmathcal%7BB%7D%20(V_%7Bt%2Cj%7D%2C%201%20-%20e%5E%7B%5Cxi_%7Bt%2Cj%7D%7D)%20#0>

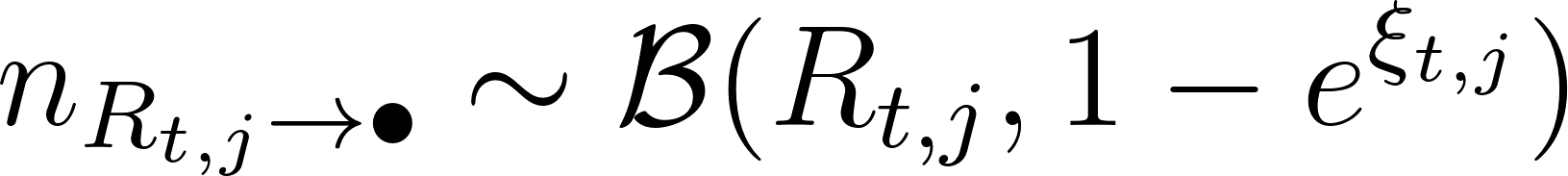
<https://www.codecogs.com/eqnedit.php?latex=%20n_%7BR_%7Bt%2Cj%7D%20%5Crightarrow%20%5Cbullet%7D%20%5Csim%20%5Cmathcal%7BB%7D%20(R_%7Bt%2Cj%7D%2C%201%20-%20e%5E%7B%5Cxi_%7Bt%2Cj%7D%7D)%20#0>

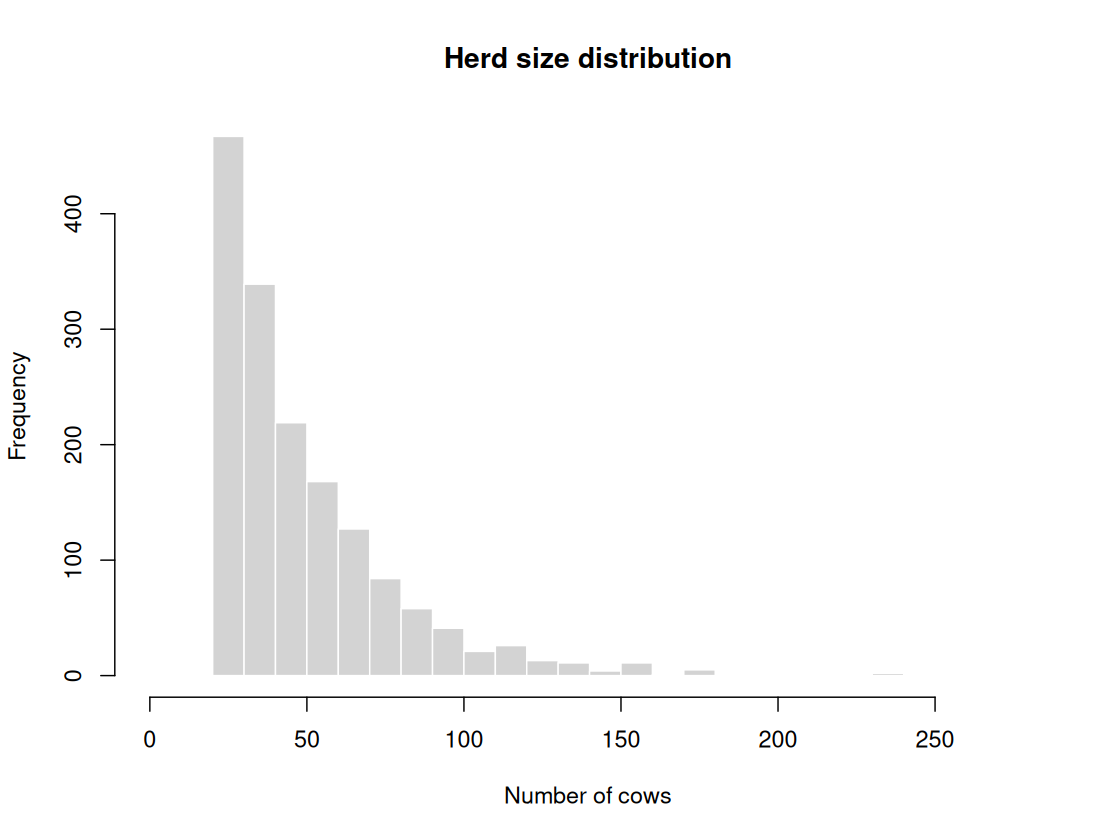

### Figure S1: Herd size distribution used in simulated LSD epidemics.

This histogram shows the distribution of herd sizes, defined as the number of cattle living in a farm. It was generated as an exponential distribution with mean 30, with an offset of 20, resulting in an average of 50 cattle per farm.

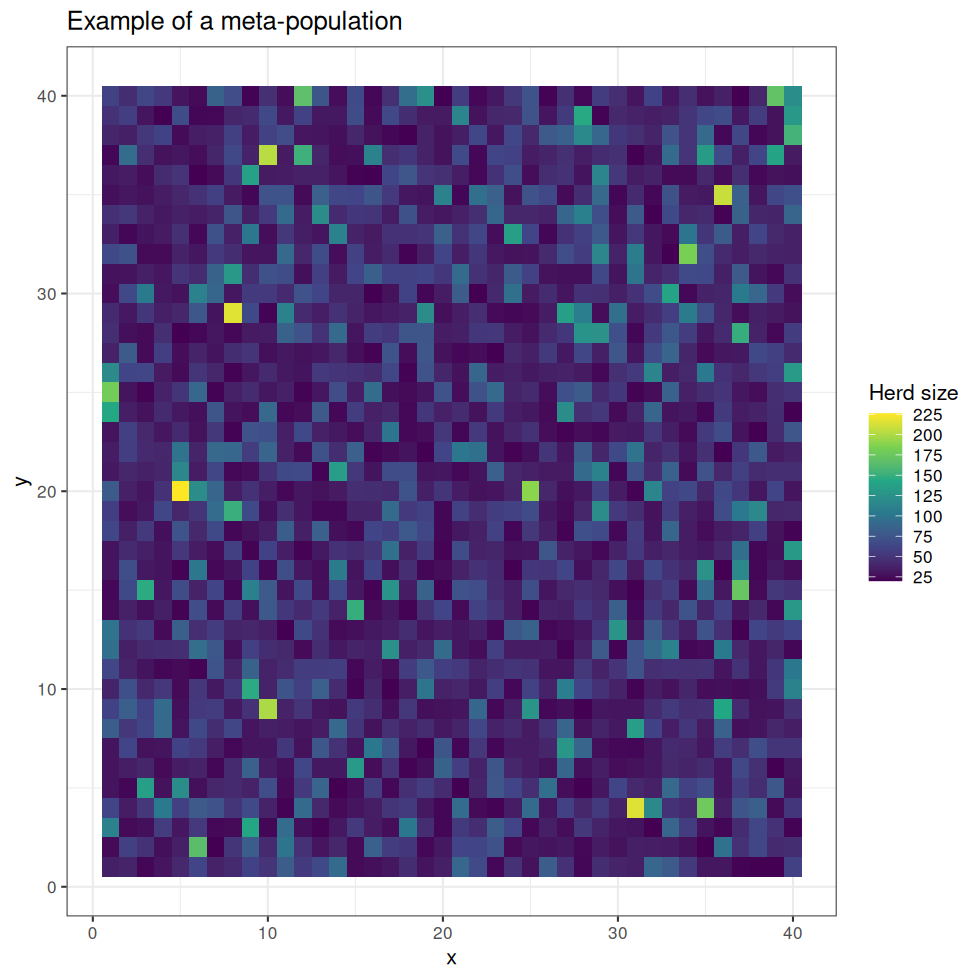

### Figure S2: Example of geographic distribution of farms.

This raster illustrates the geographic distribution of one randomly selected meta-population setting amongst the 1,000 used in our simulations. Each pixel represents a simulated farm, colored according to the size of its herd. The x and y axes represent fictitious longitudes and latitudes. Each farm is considered a neighbour to all the farms immediately right, left, up, or down from itself, so that the dispersal network (non-null off-diagonal values of
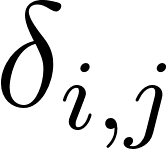
<https://www.codecogs.com/eqnedit.php?latex=%5Cdelta_%7Bi%2Cj%7D#0>) uses rook connectivity.

### Table S1: Number of LSD cases in the simulated epidemics.

This table reports, by response scenario, the means, median, 2.5% and 97.5% quantiles of the numbers of cases after a year of simulation, as well as the proportion of simulations with more than 100 cases. See Table 4 (main text) for details of the simulated scenarios.

| **scenario** | **mean** | **median** | **quantile 2.5%** | **quantile 97.5%** | **p(>100 cases)** |
| --- | --- | --- | --- | --- | --- |
| *baseline* | 25K | 25K | 1 | 42K | 0.944 |
| *cull_quarant_vacc* | 3 | 2 | 1 | 6 | 0.000 |
| *quarant_vacc* | 42 | 37 | 1 | 110 | 0.034 |
| *selcull_quarant_vacc* | 9 | 8 | 1 | 24 | 0.000 |
| *selcull_quarant_vacc_insect* | 6 | 5 | 1 | 14 | 0.000 |
| *all* | 2 | 2 | 1 | 6 | 0.000 |

### Table S2: Number of LSD deaths in the simulated epidemics.

This table reports, by response scenario, the means, median, and 2.5% and 97.5% quantiles of the numbers of deaths after a year of simulation. See Table 4 (main text) for details of the simulated scenarios.

| **scenario** | **mean** | **median** | **quantile 2.5%** | **quantile 97.5%** |
| --- | --- | --- | --- | --- |
| *baseline* | 739 | 724 | 0 | 1K |
| *cull_quarant_vacc* | 70 | 56 | 23 | 201 |
| *quarant_vacc* | 1 | 1 | 0 | 5 |
| *selcull_quarant_vacc* | 9 | 8 | 0 | 24 |
| *selcull_quarant_vacc_insect* | 6 | 5 | 0 | 14 |
| *all* | 66 | 54 | 22 | 184 |

### Table S3: Attack rates of LSD in the simulated epidemics.

This table reports, by response scenario, the means, median, and 2.5% and 97.5% quantiles of the attack rates of LSD after a year of simulation. See Table 4 (main text) for details of the simulated scenarios.

| **scenario** | **mean** | **median** | **quantile 2.5%** | **quantile 97.5%** |
| --- | --- | --- | --- | --- |
| *baseline* | 0.3196 | 0.3195 | 0 | 0.5383 |
| *cull_quarant_vacc* | 0.0000 | 0.0000 | 0 | 0.0001 |
| *quarant_vacc* | 0.0005 | 0.0005 | 0 | 0.0014 |
| *selcull_quarant_vacc* | 0.0001 | 0.0001 | 0 | 0.0003 |
| *selcull_quarant_vacc_insect* | 0.0001 | 0.0001 | 0 | 0.0002 |
| *all* | 0.0000 | 0.0000 | 0 | 0.0001 |

### Table S4: Number of farms with at least one LSD case in the simulated epidemics.

This table reports, by response scenario, the means, median, and 2.5% and 97.5% quantiles of the numbers of farms affected after a year of simulation. See Table 4 (main text) for details of the simulated scenarios.

| **scenario** | **mean** | **median** | **quantile 2.5%** | **quantile 97.5%** |
| --- | --- | --- | --- | --- |
| *baseline* | 552.55 | 550 | 1 | 920.02 |
| *cull_quarant_vacc* | 1.29 | 1 | 1 | 3.00 |
| *quarant_vacc* | 1.96 | 2 | 1 | 5.00 |
| *selcull_quarant_vacc* | 1.26 | 1 | 1 | 3.00 |
| *selcull_quarant_vacc_insect* | 1.19 | 1 | 1 | 2.00 |
| *all* | 1.23 | 1 | 1 | 3.00 |
